# C3aR1 on β cells enhances β cell function and survival

**DOI:** 10.1101/2024.11.11.622969

**Authors:** Renan Pereira de Lima, Ang Li, Ankit Gilani, James C Lo

## Abstract

Pancreatic β cell dysfunction is critical to the development of type 2 diabetes (T2D). We show that the complement receptor C3aR1 on β cells plays an essential role in maintaining β cell homeostasis, especially under the metabolic duress of obesity and T2D. Mice with β cell specific deletion of *C3ar1* have worse glucose tolerance, lower insulin levels, and decreased β cell mass. Islets from β cell specific *C3ar1* knockout (β-C3aR1 KO) mice demonstrate impaired insulin secretion. Disruption of *C3ar1* on β cells ablates the insulin secretory response to C3a, establishing a signaling axis between C3a and β cell-derived C3aR1. Markers of β cell identity were decreased while stress markers were increased in β-C3aR1 KO mice. Islets from β-C3aR1 KO also exhibit increased β cell death to lipotoxicity. Finally, we show that *C3AR1* is positively correlated with insulin secretion in human islets. These findings indicate that C3aR1 expression on β cells is necessary to maintain optimal β cell function and preserve β cell mass in T2D.

## Introduction

Type 2 diabetes (T2D) is a multi-organ disease with profound impacts on cancer, cardiovascular, kidney and retinal diseases (1, 2). Diabetes mellitus has become a pandemic afflicting over 38 million people in the USA with over 90% having T2D (3). In the US alone, the economic costs of diabetes in 2022 accounts for $306 billion in direct medical costs (4). β cell dysfunction is key to developing T2D with insufficient insulin secretion to compensate for the insulin resistance (5, 6). A hallmark of chronic T2D is pancreatic β cell failure, resulting in insulinopenia and severe hyperglycemia that requires insulin therapy(7). There are two major antihyperglycemic pharmacological therapies that target β cells in current clinical use for T2D: sulfonylureas and incretin mimetics or dipeptidyl peptidase-4 (DPP-4) inhibitors that act on the same pathway (2). Sulfonylureas have been associated with premature β cell failure(8). Whereas incretin mimetics are mostly moving towards targeting appetite suppression in the brain with GLP-1R and GIP-R co-agonists and tri-agonists with GCG-R (9, 10).

β cell defects in T2D can be attributed to β cell dysfunction and/or loss of β cell mass. Both of which can act alone or in combination to result in insufficient insulin secretion. β cell attrition from the metabolic stress of T2D can be due to cell death and trans- or de-dedifferentiation (11, 12). It is unknown if β cells that have adopted a different cellular identity can regain their β cell phenotype to secrete insulin. Strategies to augment β cell mass in T2D have yet to make it into the clinic and include inducing β cell proliferation and transplantation.

Complement proteins are expressed within the pancreatic islets and in β cells and can regulate insulin secretion (13). We had previously shown that the adipokine adipsin/complement factor D(cfd) was part of an adipose-pancreas organ cross talk that boosted glucose-stimulated insulin secretion through the C3a peptide that is generated downstream of adipsin action (14, 15). Furthermore, adipsin prevented the development of β cell failure in *db/db* mice (16). In humans, patients with T2D or worse β cell functional parameters such as HOMA-β have lower circulating adipsin levels compared to controls at earlier stage of T2D or without disease (17, 18). Higher circulating adipsin levels also associate with freedom from incident T2D in middle-aged adults (16). Adipsin may also play a protective role in protection against β cell failure in T2D in humans as patients with T2D and β cell failure have lower adipsin levels than those with T2D without β cell failure (15).

Our previous work suggested that adipsin catalyzes the formation of the C3 convertase which then generates C3a to act on the G-protein coupled receptor (GPCR) C3aR1 on the β cell to insulin secretion and protect against cell death and dedifferentiation (14, 19). However, C3aR1 is expressed on many different cell types in the islet including macrophages (13, 20). To determine the role of C3aR1 on β cells in T2D, we generated β cell specific C3aR1 knockout mice and assessed glucose handling in T2D conditions, focusing on β cell function and homeostasis. Here we now show that C3aR1 on the β cell plays a critical role in maintaining β cell function and numbers especially under metabolic stress conditions of T2D.

## Results

### C3aR1 on β cells regulates glucose homeostasis and insulin secretion

We previously found that the adipokine adipsin augments glucose stimulated insulin secretion through generation of the complement component C3a to act on islet cells (15). To directly interrogate the role of *C3ar1* on β cells in glucose homeostasis and insulin secretion, we generated β-cell specific C3aR1 knockout (β-C3aR1 KO) mice by crossing *C3ar1* floxed mice with Ins1-Cre transgenic mice. Successful deletion of *C3ar1* in pancreatic β cells from the β-C3aR1 KO mouse was confirmed by >99% decrease in *C3ar1* mRNA expression compared to Ins1-Cre heterozygous controls in sorted β cells (Figure 1A). 15- and 30-weeks old control and β-C3aR1 KO mice were challenged with a glucose tolerance test. Under the non-diabetogenic conditions of a regular diet for 15 weeks, male and female β-C3aR1 KO mice had similar glucose tolerance compared to controls (Suppl. 1A-D). However, 30 weeks old β-C3aR1 KO mice had moderately impaired glucose tolerance (Figure 1B,C) and insulin secretion (Figure 1D) compared to control mice. Female β-C3aR1 KO mice showed similar glucose tolerance but exhibited a 34% and 18% reduction in insulin secretion at 5 and 20 minutes after glucose challenge, respectively, compared to controls (Suppl. 2A,B)

**Figure 1.**
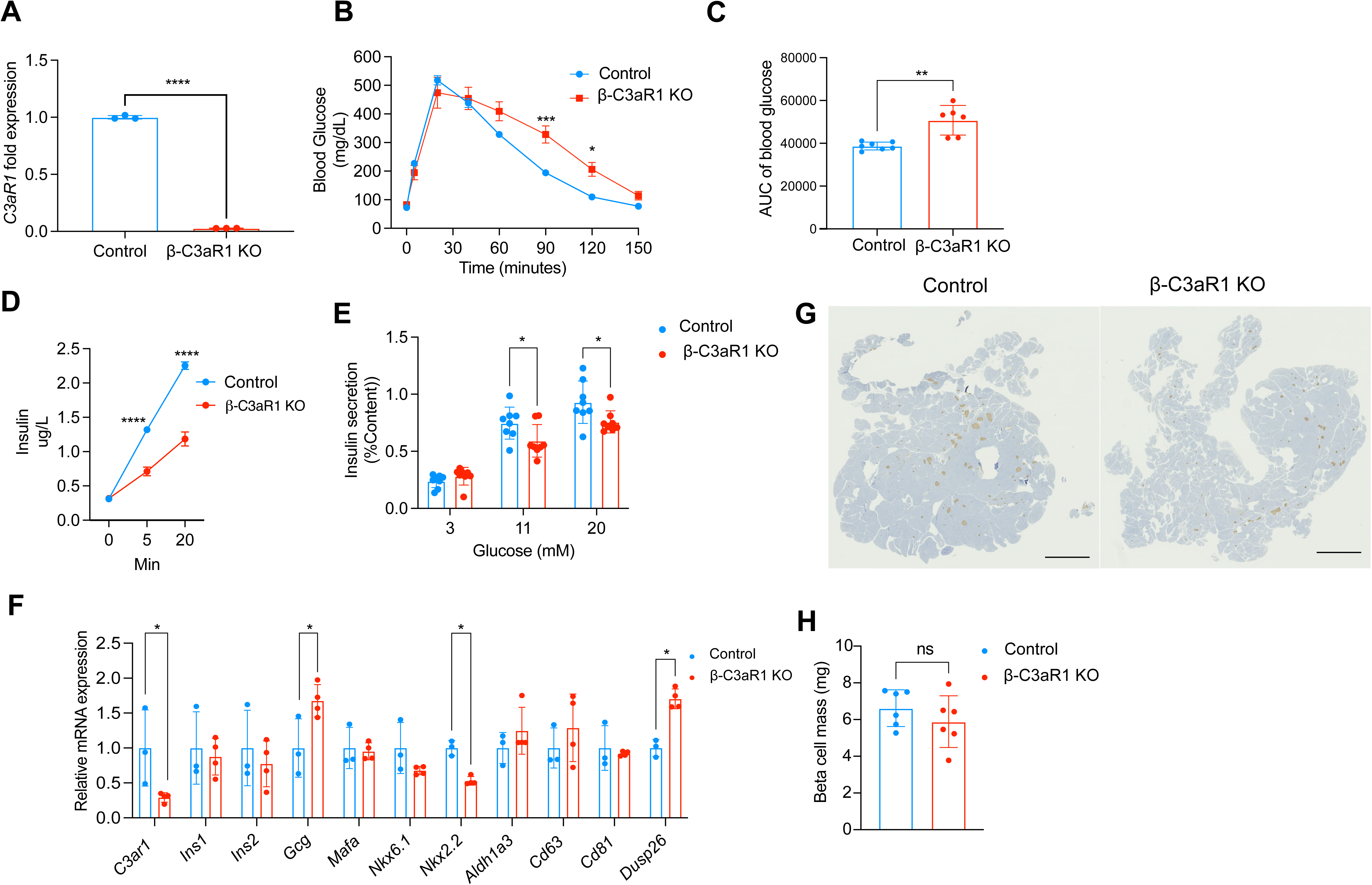
C3aR1 on β cells regulates glucose homeostasis and insulin secretion. **A**. Relative mRNA levels of *C3ar1* from sorted β cells in control and β-C3aR1 KO islets (n=3/group). **B**. Glucose tolerance test (GTT) was performed on 30-weeks-old control and β-C3aR1 KO mice (n=7/group). **C.** Glucose area under curve (GAUC) during GTT (n=6-7/group). **D.** After challenged with i.p. glucose injections plasma insulin levels were assayed (n=7/group). **E.** Glucose-stimulated insulin secretion assay on islets from 30-weeks-old control and β-C3aR1 KO mice (n=8/group). **F.** Relative gene expression of key transcription factors essential for β cell development and immature β cell (n=3/group). **G.** Representative IHC staining for insulin (brown) in pancreases of control and β-C3aR1 KO mice. Scale bars 100 μm. IHC was performed at least twice, independently, with similar results. **H.** Quantification of β cell mass in control and β-C3aR1 KO mice (n= 6/group). Data are presented as mean ± SEM. Unpaired 2-tailed t test is used for figures A, C, E, F and G and ANOVA two-way for multiple comparations is used for figures B and D. *P < 0.05, **P < 0.01, ***P < 0.001, ****P < 0.0001 vs β-C3aR1 KO mice.

To directly interrogate whether loss of C3aR1 in β cells impaired β cell function, we isolated islets from control and β-C3aR1 KO mice. Islets from β-C3aR1 KO mice exhibited approximately 20% and 18% decrease in insulin secretion following stimulation with 11 and 20 mM glucose, respectively, for both male (Figure 1E) and female mice (Suppl. 2C), recapitulating the in vivo phenotype. In addition, β-C3aR1 KO mice exhibited a marked reduction in the expression of *Nkx2.2*, a critical transcription factor essential for the development and maintenance of β cells, and elevations in *Gcg* and *Dusp26* (Figure 1F). We previously showed that C3a negatively regulates *Dusp26* expression in β cells, while palmitate enhances its expression, suggesting that C3a plays a protective role in mitigating β cell dysfunction under lipotoxicity by inhibiting *Dusp26* (16). To assess for a potential quantitative defect in β cells, we performed insulin immunohistochemistry and found no significant differences in islet morphology and β cell mass between control and β-C3aR1 KO mice (Figure 1 G,H).

### β cell derived C3aR1 is important in maintaining glucose homeostasis and insulin secretion under the metabolic stress of obesity

At 4 weeks of age, we subjected control and β-C3aR1 KO mice to high fat diet (HFD) to explore the role of C3aR1 on β cells in regulating whole-body energy and glucose homeostasis under metabolic stress. We compared the body weights of control and β-C3aR1 KO mice subjected to a HFD. After 15 weeks on a HFD, we did not observe any differences in glucose tolerance between male and female β-C3aR1 KO mice and their controls (Suppl. 3A-D). After 25 weeks on a longer-term HFD, we did not observe any significant differences in body weight between β-C3aR1 KO mice and control, in either males or females (Figure 2A; Suppl. 4A). However, after 25 weeks of HFD, male β-C3aR1 KO mice exhibited moderate impairments in glucose tolerance (Figure 2B,C). No changes were observed between female control and β-C3aR1 KO mice (Suppl. 4B,C). Control and β-C3aR1 KO mice on a HFD demonstrated similar insulin tolerance tests, suggesting that changes in insulin sensitivity were not responsible for the differences in glucose tolerance (Figure 2 D,E).

**Figure 2.**
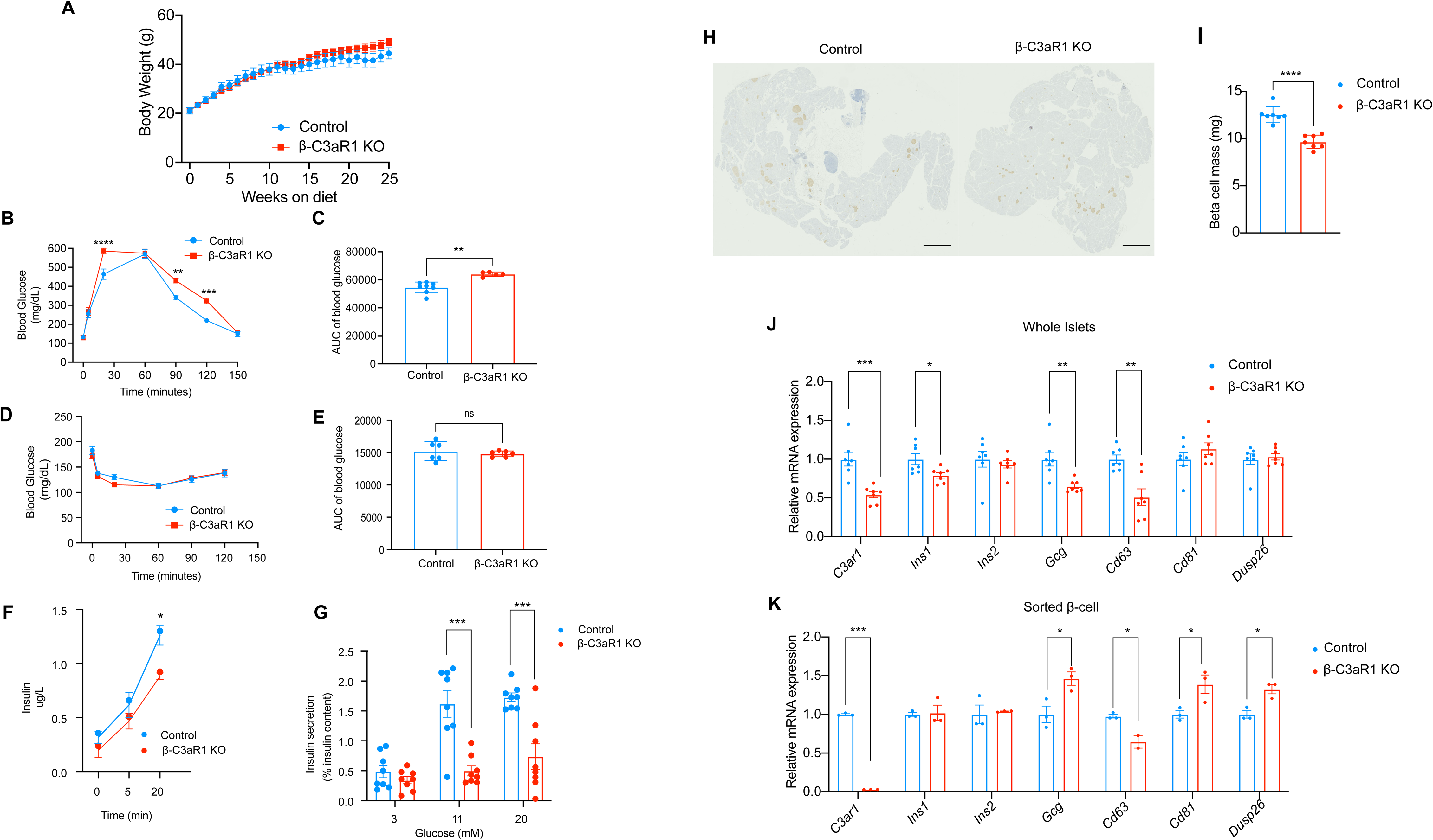
β cell derived C3aR1 is important in maintaining glucose homeostasis and insulin secretion under the metabolic stress of obesity. **A.** Body weights of control and β-C3aR1 KO male mice fed a high fat diet (HFD) for 25 weeks starting at 4 weeks (n=8/group). **B.** Glucose tolerance test (GTT) was performed on control and β-C3aR1 KO mice (n=5-8/group). **C.** Glucose area under curve (AUC) during GTT (n=5-8/group). **D**. Insulin tolerance test (ITT) was performed on control and β-C3aR1 KO mice fed a HFD for 25 weeks with measurement of blood glucose concentrations (n=6/group). **E**. Glucose area under the curve (AUC) during ITT (n=6/group). **F**. Plasma insulin levels from control and β-C3aR1 KO male mice on HFD after i.p. glucose challenge at the indicated time points (n=5-8/group). **G.** Glucose-stimulated insulin secretion assay on islets from control and β-C3aR1 KO male mice after 25 weeks on HFD (n=8/group). **H.** Representative IHC staining for insulin (brown) in pancreases of control and β-C3aR1 KO male mice fed a HFD for 25 weeks. Scale bars 100 μm. IHC was performed at least twice, independently, with similar results. **I.** Quantification of β cell mass in control and β-C3aR1 KO mice (n=7/group). **J,K**. Relative gene expression of **(J)** whole islets (n=7/group) and **(K)** sorted β cells (n=3/group) from control and β-C3aR1 KO male mice on HFD. Data are presented as mean ± SEM. Unpaired 2-tailed t test is used for figures C, E, G, H and K or ANOVA two-way for multiple comparations is used for figures A, B, D and F. *P < 0.05, **P < 0.01, ***P < 0.001, ****P < 0.0001.

To evaluate the potential role of C3aR1 on β cells in regulating insulin secretion, we measured insulin levels in control and β-C3aR1 KO mice subjected to a HFD. Male and female β-C3aR1 KO mice on a longer term HFD exhibited lower insulin levels following a glucose challenge, suggesting an important role for C3aR1 on β cells to induce insulin secretion (Figure 2F, Suppl. 4D). These findings implicated β cell insufficiency as an explanation for the exacerbated hyperglycemia in β-C3aR1 KO mice. This may result from decreased β cell mass and/or β cell dysfunction. To directly interrogate whether loss of C3aR1 in β cells impaired β cell function, we performed glucose stimulated insulin secretion (GSIS) on isolated islets. Islets from male β-C3aR1 KO mice displayed a dramatic reduction in insulin secretion in response to glucose compared to controls (Figure 2G). There was a similar phenotype in females with a moderate reduction in glucose-stimulated insulin secretion (Suppl. 4E). To evaluate for a quantitative defect in β cells, we performed insulin immunohistochemistry on pancreatic sections and found that β-C3aR1 KO mice exhibited a ∼23% reduction in β cell mass (Figure 2H,I). Collectively, these results indicate that C3aR1 is important for both the maintenance of insulin secretion and the preservation of pancreatic β cells.

To further interrogate the role of C3aR1 in β cell maintenance, we analyzed genes involved in β cell identity and function. In islets from male mice, we observed that loss of C3aR1 on β cells resulted in mild-moderate decrease in *Ins1* and halving of *Cd63*, a marker of a subset of β cells with enhanced insulin secretion (Figure 2J) (21). In sorted β cells from male mice, we found a moderate reduction in *Cd63* alongside increases in *Gcg*, *Dusp26*, and *Cd81*, the last of which is a marker of β cell stress (Figure 2L) (21, 22). In islets from female mice, *Gcg* was similarly increased in the β-C3aR1 KO group compared to controls (Suppl. 4F).

### C3aR1 on β cells is important for maintenance of β cell identity

To determine if C3aR1 on β cells plays a role in β cell identity, we analyzed the expression key transcription factors essential for β cell identity. In whole islets, we observed that loss of C3aR1 on β cells resulted in >50% reduction in *Mafa* in β-C3aR1 KO mice compared to controls (Figure 3A). *Mafa* was decreased by ∼70% and *Nkx6.1* by ∼25% in the β-C3aR1 KO group in sorted β cells (Figure 3B). *Aldh1a3* was elevated with C3aR1-deficiency in both islets and sorted β cells (Figure 3A,B). Female mice also demonstrated marked reductions in islet *Mafa* and *Nxk6.1* in the β-C3aR1 KO group compared to controls (Suppl. 4F).

**Figure 3.**
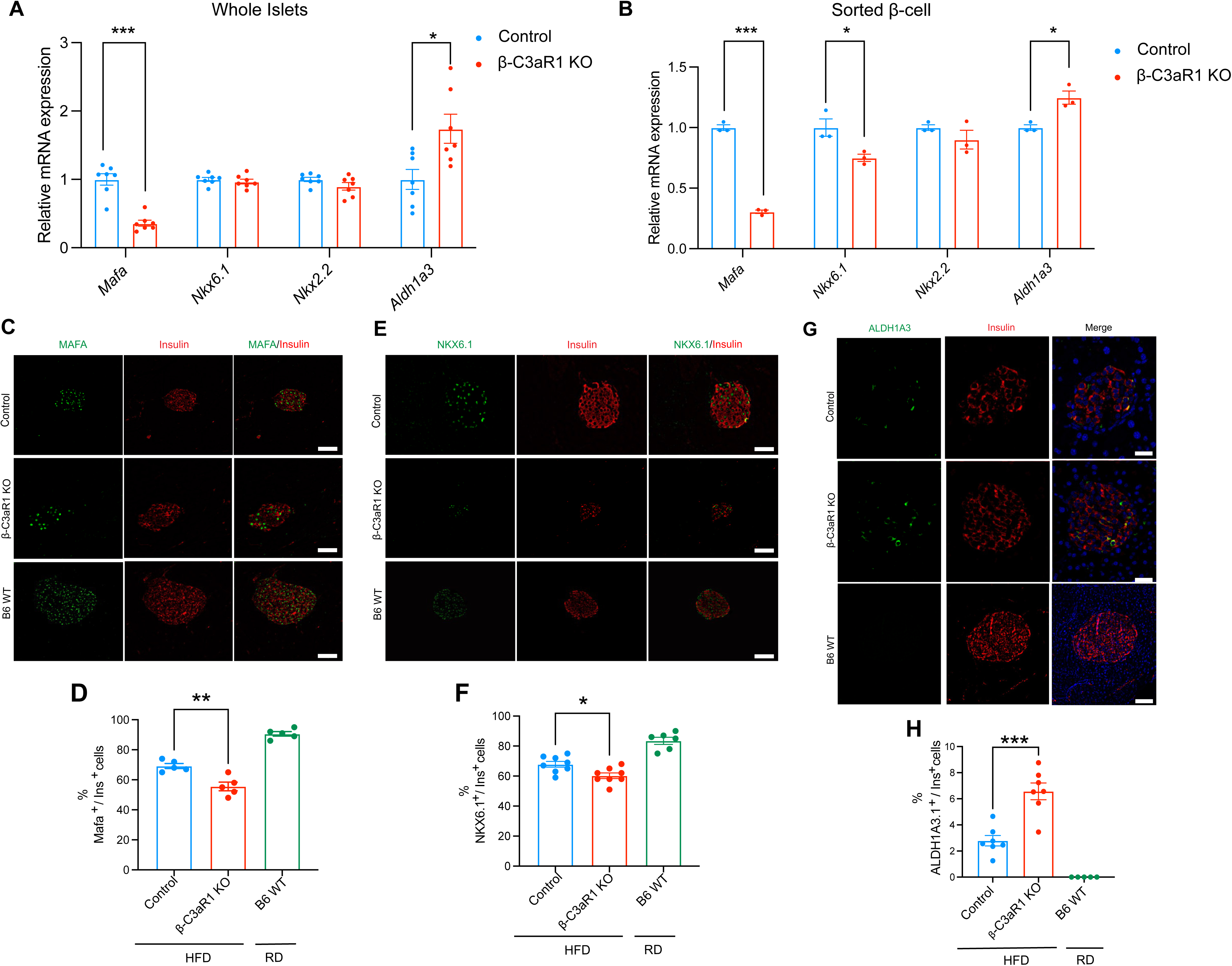
C3aR1 on β cells is important for maintenance of β cell identity. **A,B.** Gene expression of key transcription factors essential for β cell identity in whole islets (**A)** (n=7/group) and sorted β cells (**B)** (n=3/group) from control and β-C3aR1 KO male mice fed a high fat diet (HFD) for 25 weeks**. C,D.** Representative images of MAFA and insulin immunofluorescence (IF) in B6 WT mice on regular diet (RD) and control and β-C3aR1 KO mice on HFD with quantification of MAFA+ β cells **(D)** (n=5/group). **E,F.** Representative images of NKX6.1 and insulin IF in B6 WT mice on RD and control and β-C3aR1 KO mice on HFD with quantification of NKX6.1+ β cells (**F)** (n=6-8/group). **G,H.** Representative images of ALDH1A3 and insulin IF in B6 WT mice on RD and control and β-C3aR1 KO mice on HFD with quantification of ALDH1A3^hi^ β cells **(H)** (n=5-7/group). Scale bars 100 μm. Data are presented as mean ± SEM. Unpaired 2-tailed t test is used for comparison between two groups. *P < 0.05, **P < 0.01, ***P < 0.001 vs β-C3aR1 KO mice.

To investigate whether β cell C3aR1 has an effect on the maintenance of β cell transcriptional identity at the single cell level, we performed immunostaining for MAFA, NKX6.1 and NKX2.2. Deletion of C3aR1 on β cells led to a 20% decrease in β cells expressing MAFA, and ∼10% decrease in NKX6.1 without significant changes for NKX2.2 (Figure 3C-F, Suppl. 5A,B). We also performed immunostaining for glucagon and aldehyde dehydrogenase 1a3 (ALDH1A3), which is molecular marker of failing or dedifferentiated β cells. β cell specific loss of C3aR1 doubled the number of ALDH1A3^hi^ β cells, but had no significant impact on the number of glucagon positive cells on islets (Figure 4G,H and Suppl. 5C-D). Our results suggest that C3aR1 protects against β cell failure by maintaining β cell transcriptional identity.

**Figure 4.**
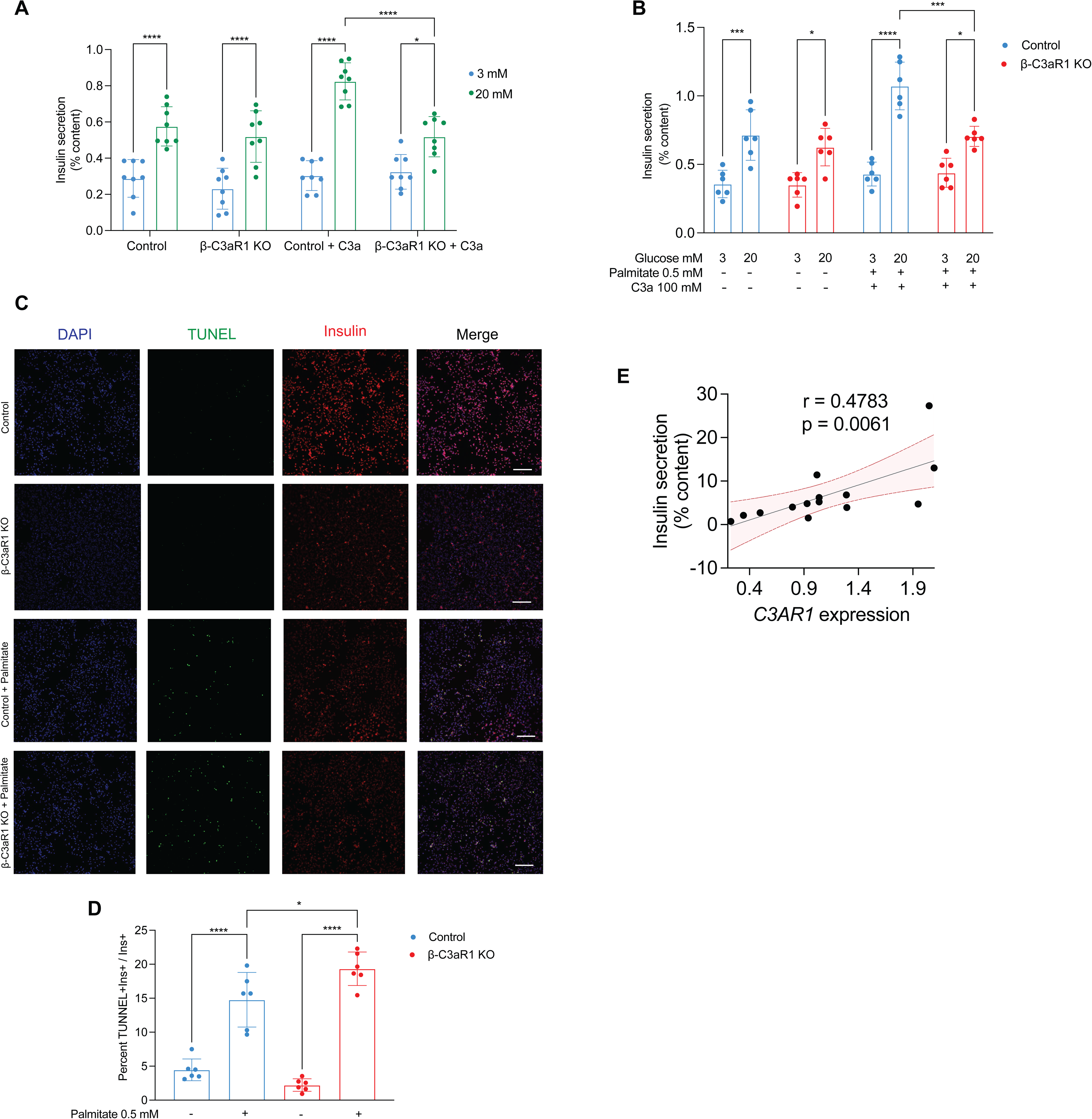
β Cell C3aR1 in C3a-mediated Insulin Secretion, Cell Death, and Human Islet Function. **A.** Islets from control and β-C3aR1 KO mice on a regular diet were subjected to a GSIS assay with recombinant C3a (100 nM) at the indicated concentrations of glucose (n=8/group). **B.** Islets were treated with vehicle (Veh) or palmitate (Pa) (0.5 mM) to a GSIS assay with recombinant C3a (100 nM) at the indicated concentrations of glucose (n=6/group). Dispersed islets were treated with Veh or Pa (0.5 mM) for 18 hours. **C.** Representative images are presented of IF staining for insulin and TUNEL assay in dispersed islets Images are representative of two independent experiments. **D.** Quantification of TUNEL+ beta cells as determined by IF (n=6/group). Scale bars, 100 μm. **E.** Correlation between human islet *C3AR1* expression and insulin secretion to 16.7mM glucose (n=14 donors). Data are presented as mean ± SEM. ANOVA one-way for multiple comparations is used for figures A, B. and D. *P < 0.05, ***P < 0.001, ****P < 0.0001 vs β-C3aR1 KO mice. Linear regression with Pearson correlation coefficient analysis (bars indicate the 95% confidence intervals).

### β Cell C3aR1 in C3a-mediated Insulin Secretion, Cell Death, and Human Islet Function

We had previously shown that C3a enhances GSIS on islets (15). It was unclear if this occurs through a direct interaction with C3aR1 on β cells or if another islet cell type expressing C3aR1 may be involved. To directly interrogate the requirement of β cell C3aR1 on the effect of C3a on GSIS, islets from control and β-C3aR1 KO mice fed a RD was subjected to a GSIS assay in the presence of C3a. Acute administration of C3a increased insulin secretion by 40% under high, but not low glucose conditions in control islets (Figure 4A). However, in islets from β-C3aR1 KO mice, C3a did not further enhance insulin secretion, suggesting that C3aR1 on the β cell is essential for C3a to augment insulin secretion (Figure 4A). Next, we assessed the β cell response to lipotoxocity and its response to C3a for insulin secretion. Islets from control and β-C3aR1 KO mice on a RD were exposed to palmitate for 18 hours. While control islets robustly responded to C3a even after palmitate treatment, islets from the β-C3aR1 KO group failed to respond (Figure 4B). These results suggest that C3a stimulation of C3aR1 on the β cell can augment GSIS under acute lipotoxicity conditions.

We had also shown that C3a could protect against palmitate-induced cell death of islet cells (16). It was not known if C3aR1 expression on β cells is a prosurvival factor. Thus, we used the β-C3aR1 KO model to test whether C3aR1 on β cells could play a role in protecting β cells from death. We found that ablation of C3aR1 on β cells resulted in a 30% increase in palmitate-induced β cell death, highlighting the crucial role of C3aR1 on β cells in survival from the metabolic stress of diabetes (Figure 4C,D).

Finally, we assessed the potential for translational relevance for our findings of C3aR1 on β cells to human diabetes and β cell function. We analyzed the relationship between *C3AR1* expression and insulin release in human pancreatic islets. Our results revealed that *C3AR1* positively correlates with insulin secretion in islets from non-diabetic human donors, suggesting that C3aR1 may play a role in modulating insulin secretion in human islets (Figure 4E). These findings provide new insights into the function of C3aR1 on β cells and the role C3aR1 plays in the complex regulation of pancreatic β cell function under normal physiologic and pathological conditions of T2D.

## Discussion

Our study demonstrates that C3aR1 on β cells plays an important non-redundant role in insulin secretion and preservation of β cell mass. The effects with loss of C3aR1 seen in this study occur with older age or longer exposure to diet-induced obesity. C3aR1 on the β cell protects against cell death. We have shown this in the context of lipootoxicity with palmitate, which is a commonly used trigger in experimental models. Mice lacking C3aR1 specifically on β cells also demonstrate loss of core β cell transcription factors that are key to maintaining β cell identity. Likewise, β cell loss of C3aR1 leads to increased levels of stress markers such as *Cd81*, *Aldh1a3*, and *Dusp26*. The downstream C3aR1 signaling pathways in β cells that lead to the prosurvival and enhanced insulin secretion phenotype still remain to be determined (16). One major candidate that arose was DUSP26, a dual specificity phosphatase that was decreased by adipsin and C3a (16). It is not clear how much DUSP26 contributes to the protection from cell death and dedifferentiation conferred by C3aR1. Future studies elucidating the pathophysiological role of DUSP26 in β cells should address this.

We demonstrated important roles for C3a on β cell action in this and prior studies (15, 16). What still remains unknown is the source of C3a. Does C3a derive locally or from the circulation from C3? Since C3a acts as an anaphylatoxin and can be deactivated to C3a-desArg by carboxypeptidases, systemic levels of C3a generally are not high. β cells can also make C3 though they generally are thought to lack the machinery to process it to C3a (23). Rather intracellular C3 in β cells have been shown to protect against stress and death from cytokines (23, 24). It is possible that another islet cell type such as the macrophage may locally secrete adipsin. There may be more than one beneficial signal from C3 and its products to β cell health and protection against T2D.

This study shows a positive correlation with *C3AR1* in human islets with glucose-stimulated insulin secretion. It is tempting to speculate that lower C3aR1 on human β cells would lead to decreased insulin secretion that would eventually cause β cell dysfunction and T2D. We do not formally know if C3aR1 declines in β cells in humans with T2D. Or that loss of the receptor or abrogated C3aR1 signaling in human β cells is part of the pathophysiology of T2D in humans or potentially in a subset of patients (25). Even so, a pharmacologic agent targeting C3aR1 on β cells could still be effective treatment for T2D. The potential issues with drugging C3aR1 are that its ligand C3a is an anaphylatoxin and is rapidly inactivated in the serum by carboxypeptidases. However, with biased GPCR agonists it might be possible to separate out the anaphylatoxin properties from the protective β cell effects. Another future possibility is to direct the ligand to the β cell without off-cell effects like the mast cell. Other avenues include assessing what is downstream of the C3aR1 in β cells. For example, inhibition of the phosphatase DUSP26 could be a viable therapeutic option without the untoward side effects of systemic C3aR1 agonism.

There are some limitations to our study. While we show an essential role for C3aR1 on β cells in glucose-stimulated insulin secretion, this does not formally rule out the possibility that other cell types expressing C3aR1 in the islets or elsewhere could have an indirect impact on the β cells. The direct and potential indirect effects of C3a and C3aR1 are not mutually exclusive. As stated earlier, the human *C3AR1* islet data with insulin secretion is a correlation. Studies to knockout/knockdown *C3AR1* in human β cells would be needed to show causality. Our studies were performed in the B6/J strain of mice. The generalizability to other strains of mice is not known. This could be important as immune and obesity varies depending on the genetic background.

## Material and Methods

### Animals

*C3ar1* flox/flox mice were on the C57BL/6J background as described (26). Ins1-Cre transgenic mice were purchased from The Jackson Laboratory (strain 26801). Ins1-cre heterozygous mice were used in the experiments as controls. All mice were maintained in plastic cages under a 12-hour light/12-hour dark cycle at constant temperature (22°C) with free access to water and food. Mice were fed a regular chow diet (RD; 5053, LabDiet) or a 60% high fat diet (HFD; D12492i, Research Diets) after weaning.

### Blood chemistry and serum insulin analysis

Mice were fasted for 16 hours prior to the glucose tolerance tests and injected intraperitoneally with syringe-filtered D-glucose solution (2g/kg). For insulin tolerance test, mice were fasted for 6 hours and injected with 0.5 mIU/kg insulin. Blood glucose levels were assayed by commercial glucometer (OneTouch) by tail vein blood samples. Tail vein blood was collected into tubes, centrifuged at 2000xg at 4°C, and plasma insulin levels were determined by ELISA using a standard curve (Mercodia).

### Pancreatic islet isolation and culture

Mouse pancreatic islets were isolated by perfusion of the pancreases with CIZyme (VitaCyte) through the common hepatic duct as previously described (16). Islets were then hand-picked and processed for different applications. For treatment with palmitate and/or recombinant C3a, an equal number of whole handpicked islets were cultured in 3mM glucose and 0.5 mM palmitate or vehicle and then harvested. For flow cytometry and FACS sorted islet cells, dissociation was performed with 0.05% trypsin.

For static incubation assays, islets were handpicked and transferred into basal Krebs buffer containing 3 mM glucose. After 1 hour of starvation, islets were transferred into Krebs solution containing 11 or 20 mM for 1 hour at 37°C. Supernatants were centrifuged and collected. The pellet was used to determine intracellular insulin content. Insulin secretion was measured with Insulin ELISA kit (Mercodia).

### Flow cytometry

β cells were gated and purified based on granularity and FAD/FMN (FITC) autofluorescence. FAD and NAD(P)H levels were assessed by measuring autofluorescence at and excitation wavelength of 488 and 350 nm respectively using a Sony MA-900 sorter as described (21, 27).

### TUNEL assay

For determination of apoptosis in dispersed islets, a TUNEL assay (Promega) was performed according to the manufacturer’s instructions. TUNEL+ nuclei in insulin-positive staining was determined with Fiji/ImageJ (NIH). For determination of cell viability, 50,000 INS-1 cells per well were seeded in a 24-well plate and subjected to palmitate treatment and C3a (100 mM) for 18 hours.

### RNA extraction and qPCR analysis

RNA isolation was performed using the RNeasy Micro and Mini kits (Qiagen). cDNA was synthesized through reverse transcription using a cDNA synthesis kit (Thermo). cDNA was analyzed by real-time PCR using specific gene primers and a SYBR Green Master Mix (Quanta). Primer sequences are shown in Supplementary Table 1.

### IHC and IF analyses

For histological studies of islets, pancreases were dissected, laid flat for paraffin embedding and fixed in 10% neutral-buffered formalin (VWR) overnight at 4 °C. Tissues were then transferred to 70% ethanol and subsequently embedded in paraffin and sectioned at 5 μm thickness. Sections were dewaxed, and antigen retrieval was performed using 10 mM sodium citrate buffer (pH 6.0) at boiling temperature for 14 min. For insulin IHC to determine beta cell mass, anti-insulin (Dako, IR002) antibody was incubated overnight at 4° C, followed by incubation with corresponding biotinylated secondary antibodies. Signals were developed using 3,3′-diaminobenzidine (DAB). Sections were then counterstained with hematoxylin. For IF staining, appropriate primary antibodies were incubated overnight at 4° C, followed by incubation with Alexa-Fluor conjugated secondary antibodies (Thermo). For MAFA (Bethyl, IHC-00352), NKX6.1 (DSHB, F55A12), NKX2.2 (Atlas Antibodies, HPA003468), ALDH1A3 (Novus, NBP2-15339) and Glucagon (Phoenix, H-028-05) stainings, Tyramide Signal Amplification (TSA; Thermo) was used. To calculate beta cell mass, whole sections stained with insulin and developed with DAB were scanned using a Zeiss Axioscan7 whole-slide scanner. The fraction of insulin-positive areas compared to total pancreatic tissue area (hematoxylin) was determined with Fiji/ImageJ (NIH). Beta cell mass and IF analyses were determined as previously described (16).

### Human Samples

Frozen human pancreatic islets were obtained through the Integrated Islet Distribution Program (IIDP) at City of Hope (Duarte, CA) (https://iidp.coh.org) (Suppl. Table 2). The use of human islets was approved by the IIDP as previously described (21).

### Statistical analysis

Data are presented as mean ± s.e.m. Data are derived from multiple experiments unless stated otherwise. If not mentioned otherwise in the figure legend, statistical significance is indicated by *P < 0.05, **P < 0.01, ***P < 0.001 and ****P < 0.0001. Statistical analysis was carried out using unpaired, two-tailed t-test or two-way ANOVA. GraphPad Prism 10 was used for statistical analysis. Pearson analysis was used for correlations with bars indicating the 95% confidence intervals.

## Author contributions

R.P.L and J.C.L designed the study and wrote the manuscript with input from all authors. R.PL, A.L, M.R, A.G performed and analyzed the animal experiments. R.P.L and A.L developed and analyzed the in vitro experiments. JCL conceived and supervised the study.

---

**Supplementary Table 1.**
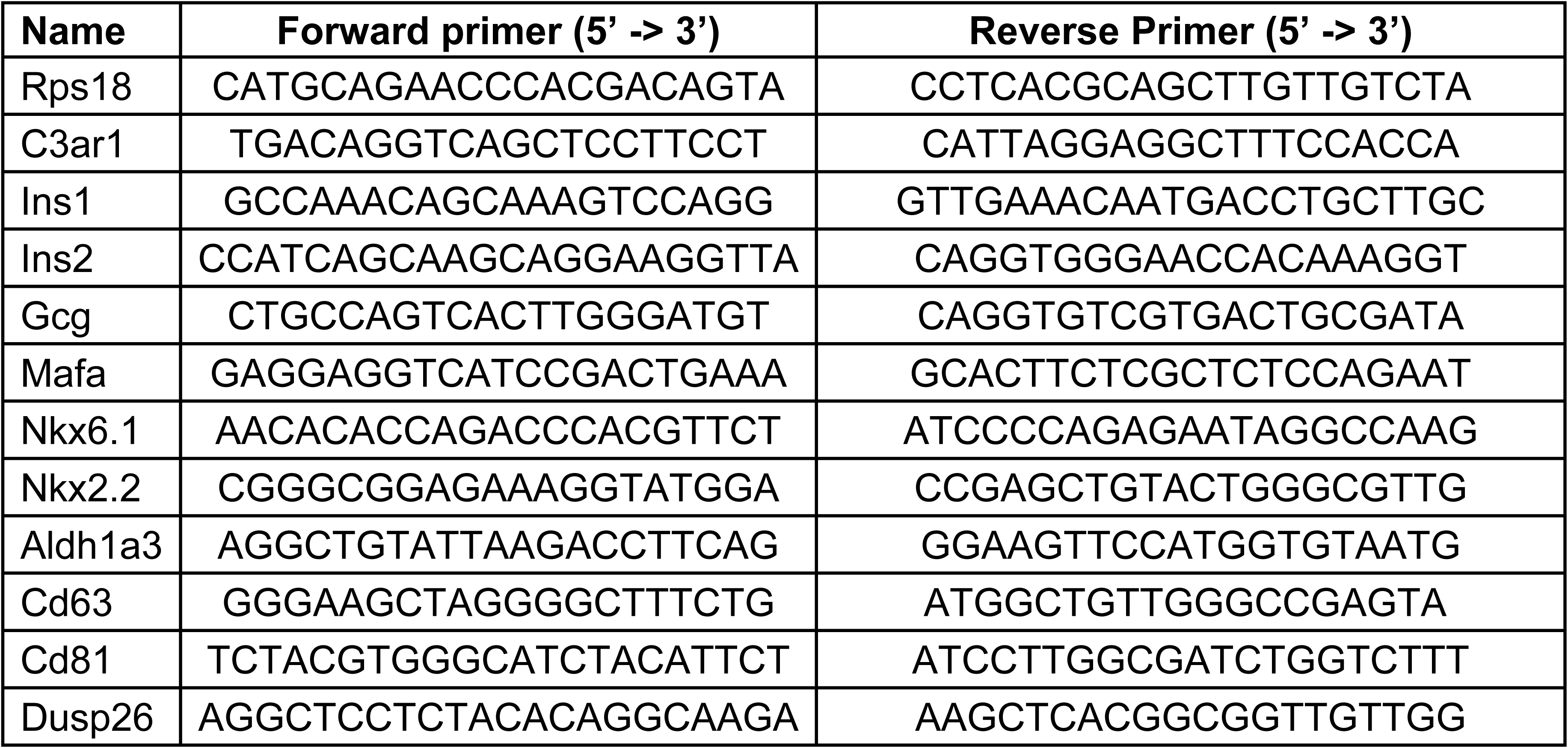
qPCR primers.

**Supplementary Table 2.**
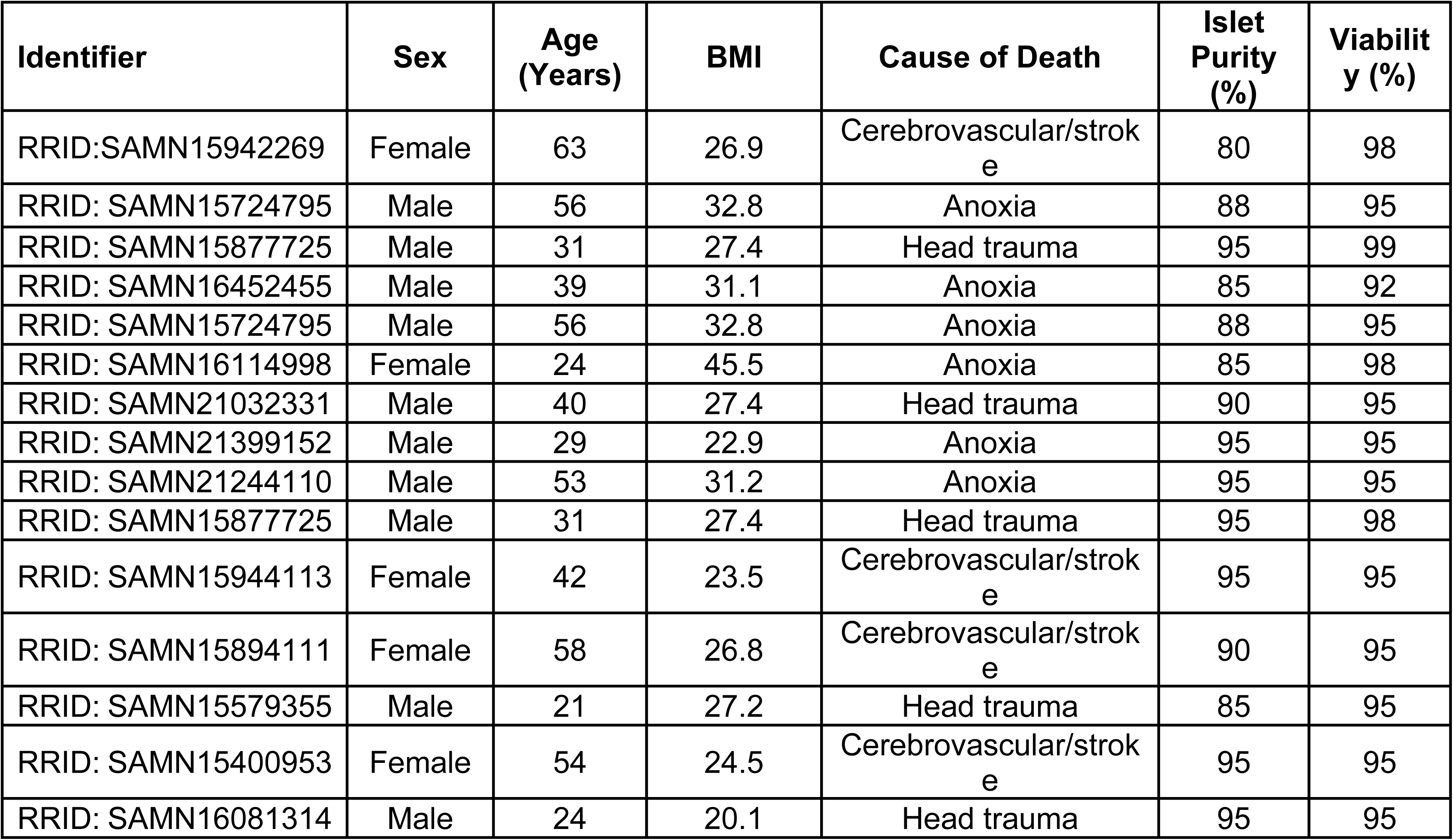
Donor characteristics for human islets.

**Supplemental Figure 1.**
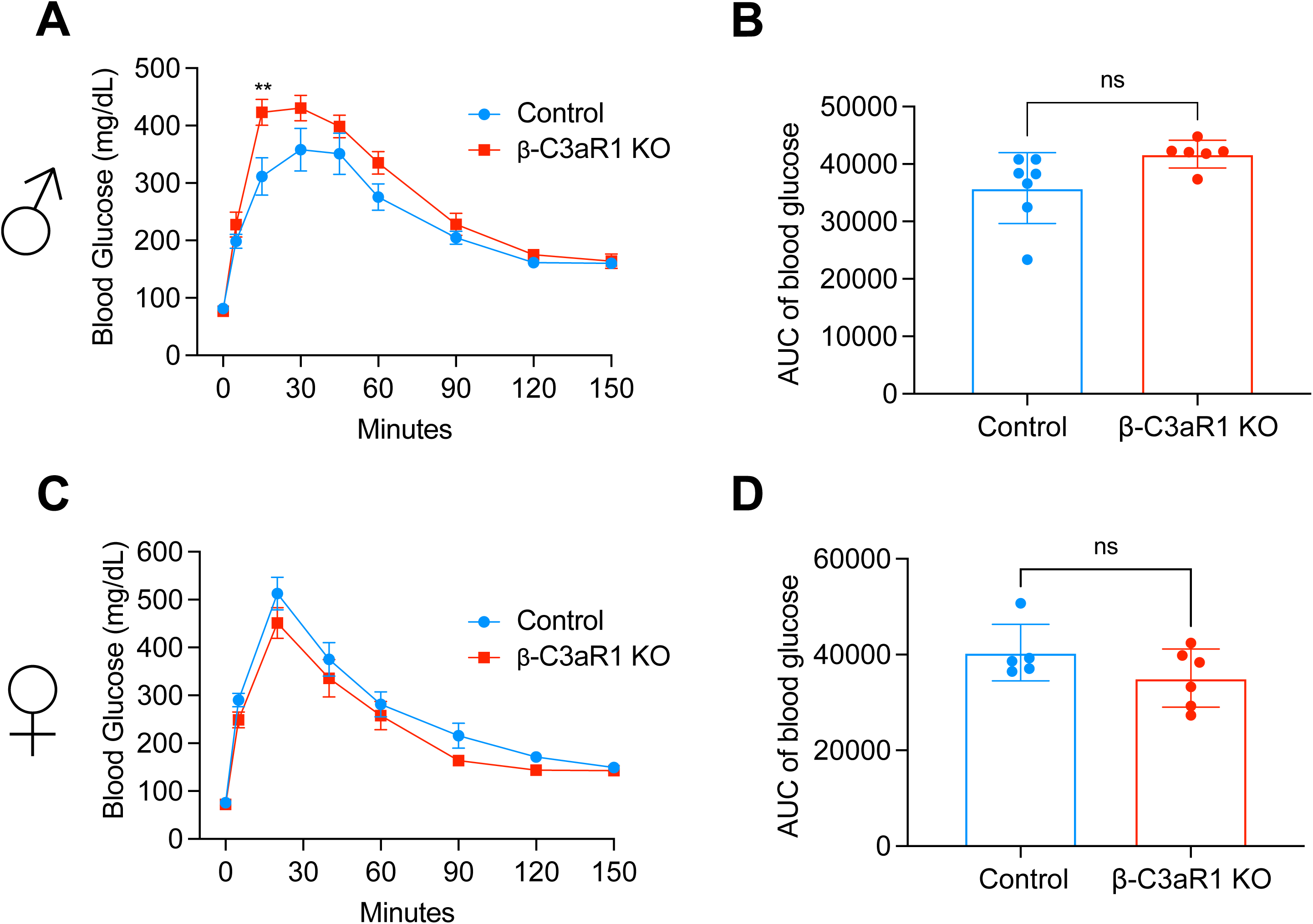
Glucose tolerance in β cell Specific C3ar1 KO male and female mice on a regular diet at 15 Weeks. **A.** Glucose tolerance test (GTT) was performed on 15 weeks-old control and β-C3aR1 KO male mice (n=6-7/group). **B.** Glucose area under curve (GAUC) during GTT (n=6-7/group). **C.** Glucose tolerance test (GTT) was performed on 15 weeks-old control and β-C3aR1 KO female mice (n=5-6/group). **D.** Glucose area under curve (GAUC) during GTT (n=5-6/group). Data are presented as mean ± SEM. Unpaired 2-tailed t test is used for figures B and D or ANOVA two-way for multiple comparations is used for figures A and C. **P < 0.01 vs β-C3aR1 KO mice.

**Supplemental Figure 2.**
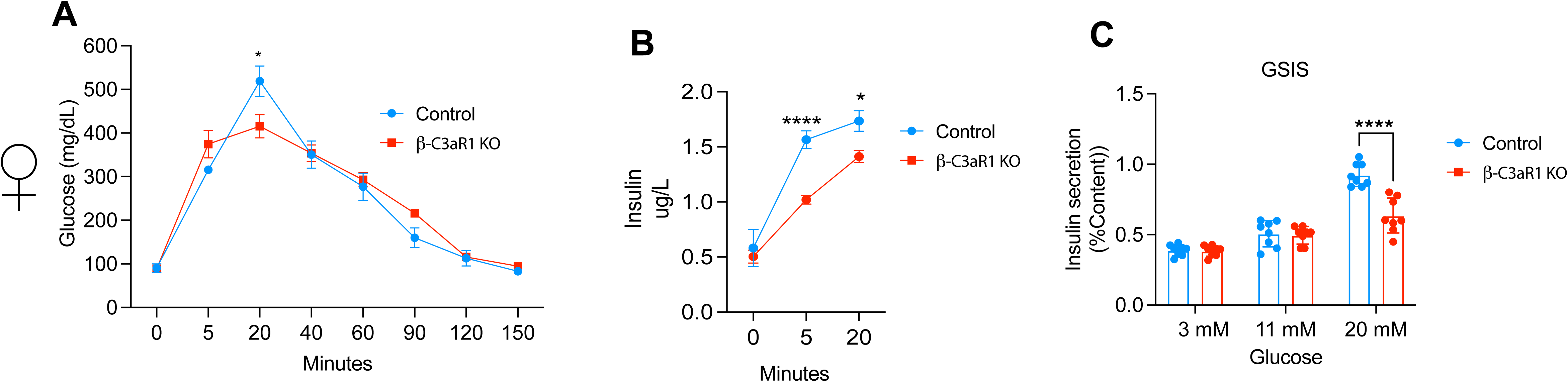
β cell C3aR1 plays a role in insulin secretion in female mice on a regular diet model. **A.** Glucose tolerance test (GTT) was performed on 30 weeks-old control and β-C3aR1 KO female mice (n=5-7/group). **B.** Glucose area under curve (AUC) during GTT (n=5-7/group). **C.** Glucose-stimulated insulin secretion assay on islets from 30-weeks-old control and β-C3aR1 KO female mice (n=8/group). Data are presented as mean ± SEM. ANOVA two-way for multiple comparations is used for figures A and B or Unpaired 2-tailed t test is used for comparison between two groups for figure C. *P < 0.05, ****P < 0.0001 vs β-C3aR1 KO mice.

**Supplemental Figure 3.**
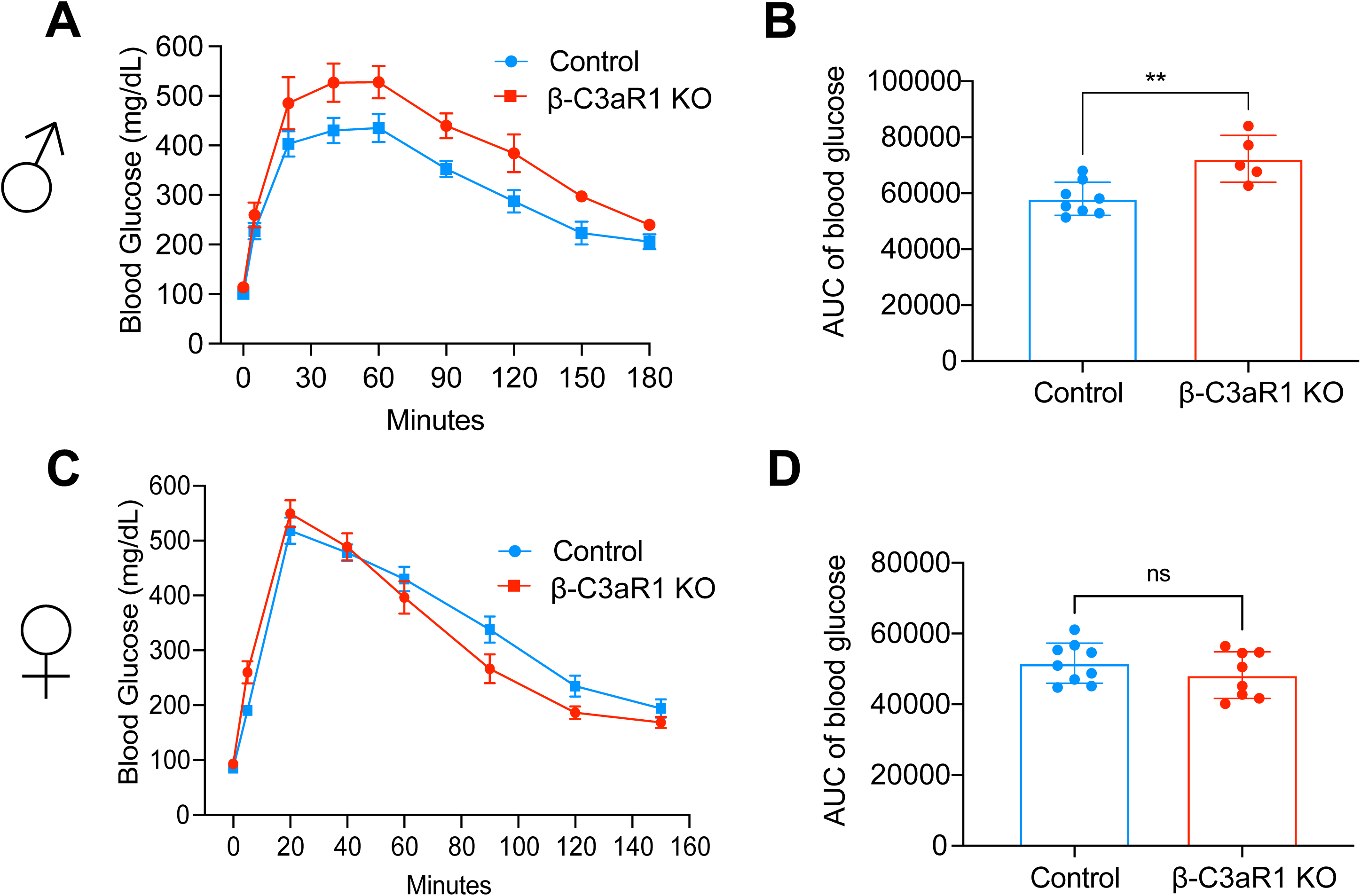
β cell C3aR1 does not play a role in insulin secretion in male and female mice on a short-term high fat diet model. A-D. Glucose tolerance test (GTT) was performed in control and β-C3aR1 KO male **(A,B)** (n=5-8/group) and female **(C,D)** (n=8-9/group) mice. **B, D.** Glucose area under curve (AUC) during GTT. Control and β-C3aR1 KO male and female mice were fed a high fat diet (HFD) for 15 weeks starting at 4 weeks of age. Data are presented as mean ± SEM. ANOVA two-way for multiple comparations is used for figures A and C or Unpaired 2-tailed t test is used for comparison between two groups for figure B and D. **P < 0.01 vs β-C3aR1 KO mice.

**Supplemental Figure 4.**
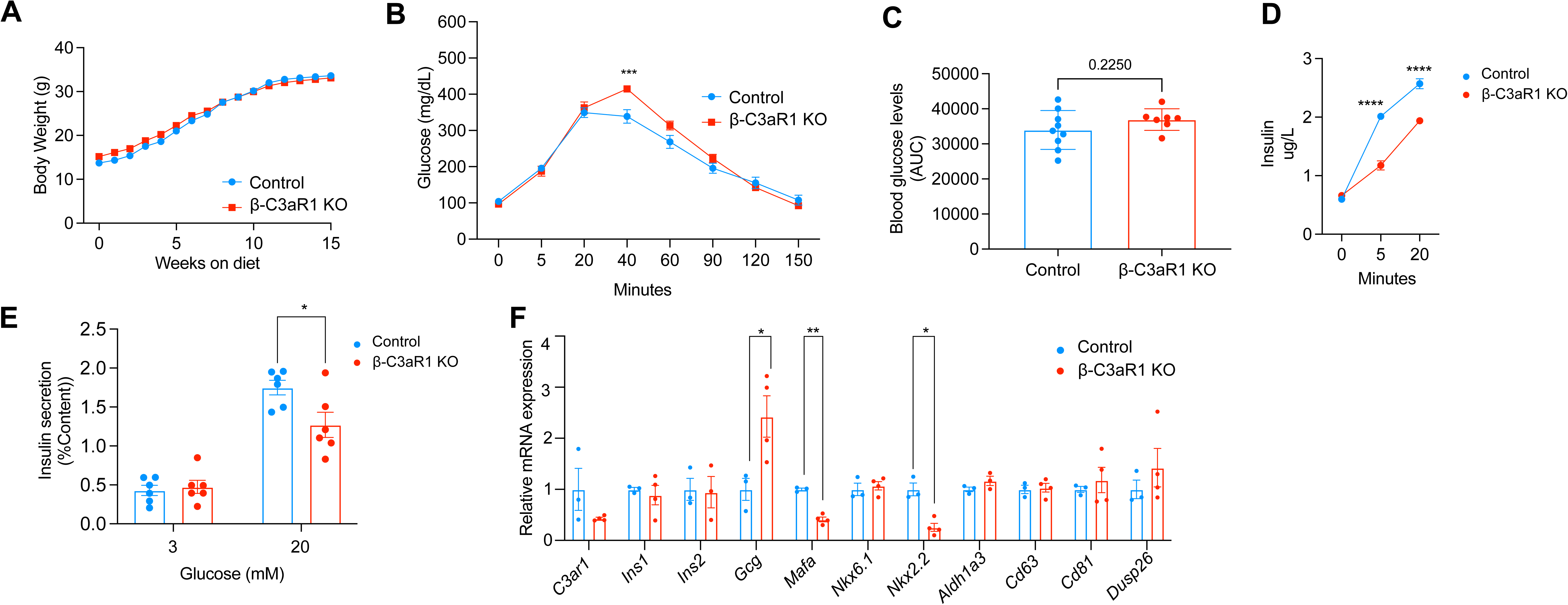
β cell C3aR1 plays a role in insulin secretion in female mice on a long-term high fat diet model. **A.** Body weights of control and β-C3aR1 KO female mice were fed a high fat diet (HFD) starting at 4 weeks of age (n=7/group). **B.** Glucose tolerance test (GTT) was performed on control and β-C3aR1 KO mice (n=7-9/group). **C.** Glucose area under curve (GAUC) during GTT (n=7-9/group). **D.** After challenged with i.p. glucose injections plasma insulin levels were assayed (n=6-9/group). **E.** Glucose-stimulated insulin secretion assay on islets from 25 weeks HFD control and β-C3aR1 KO female mice (n=6/group). **F.** Relative gene expression of key transcription factor essential for β cell development and immature β cell in whole islets (n=3/group). Data are presented as mean ± SEM. ANOVA two-way for multiple comparations is used for figures A, B and D or Unpaired 2-tailed t test is used for comparison between two groups for figure C,E and F. *P < 0.05 **P < 0.01, ***P < 0.001, ****P < 0.0001 vs β-C3aR1 KO mice.

**Supplemental Figure 5.**
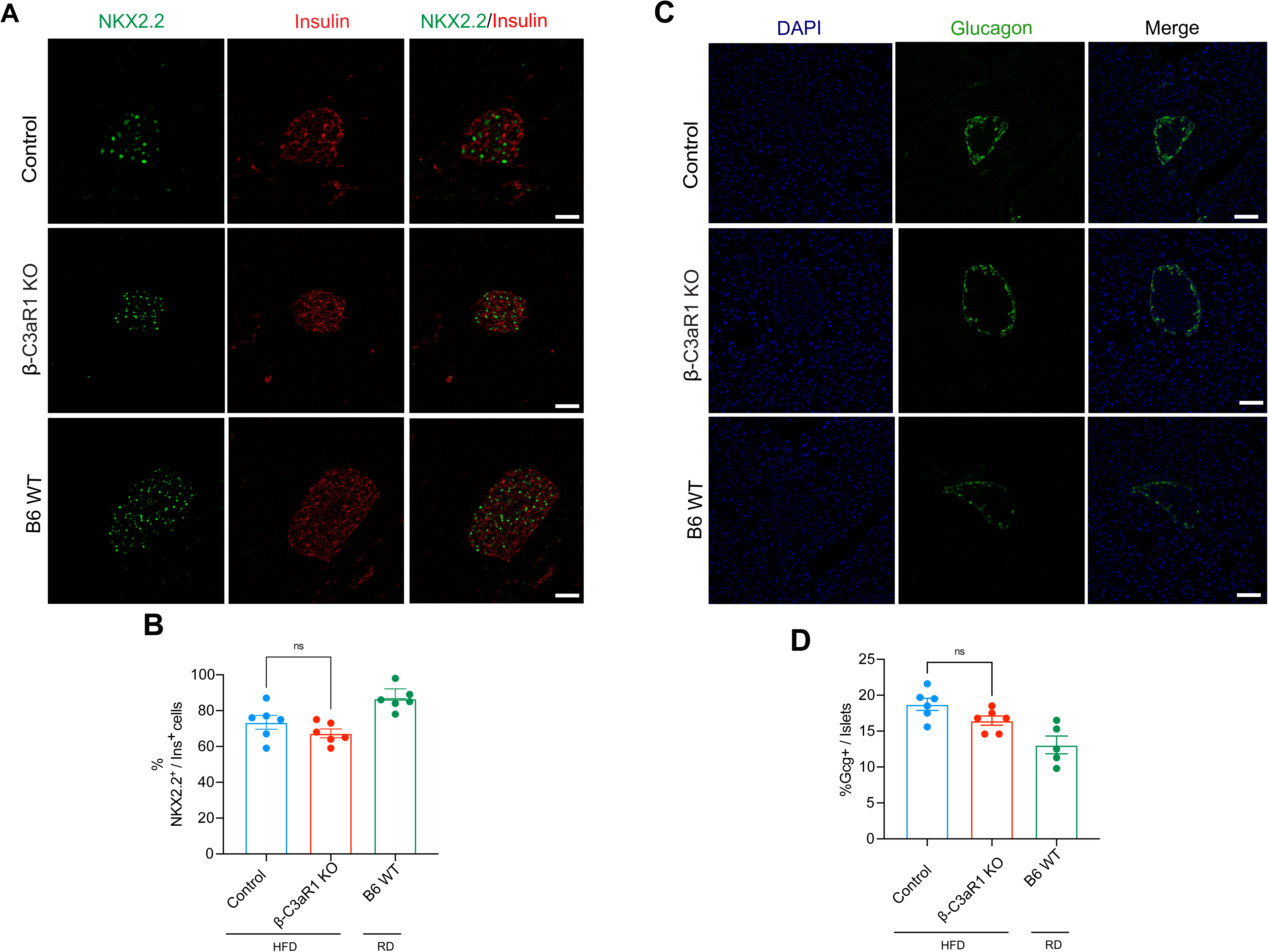
Loss of β-cell C3aR1 does not affect NKX2.2 and Glucagon expression in a long-term high-fat diet model. Control and β-C3aR1 KO mice were fed a high fat diet (HFD) for 25 weeks starting at 4 weeks. **A.** Representative images of NKX2.2 and insulin IF in B6 WT mice, control and β-C3aR1 KO mice with quantification of NKX2.2+ beta cells **(D)** (n=6/group). **E.** Representative images of glucagon and DAPI IF in B6 WT mice, control and β-C3aR1 KO mice with quantification of glucagon+/islets (**F)** (n=5-6/group). Scale bars 100 μm. Data are presented as mean ± SEM.

